# Evidence that the SARS-CoV-2 S protein undergoes a conformational change at the Golgi Complex that leads to the formation of virus neutralising antibody binding epitopes in the S1 protein subunit

**DOI:** 10.1101/2024.05.01.591558

**Authors:** Yanjun Wu, Soak Kuan Lai, Conrad En-Zuo Chan, Boon Huan Tan, Richard J Sugrue

## Abstract

Recombinant SARS-CoV-2 S protein expression was examined in Vero cells by imaging using the human monoclonal antibody panel (PD4, PD5, sc23, and sc29). The PD4 and sc29 antibodies recognised conformational specific epitopes in the S2 protein subunit at the Endoplasmic reticulum and Golgi complex. While PD5 and sc23 detected conformationally specific epitopes in the S1 protein subunit at the Golgi complex, only PD5 recognised the receptor binding domain (RBD). A comparison of the staining patterns of PD5 with non-conformationally specific antibodies that recognises the S1 subunit and RBD suggested the PD5 recognised a conformational structure within the S1 protein subunit. Our data suggests the antibody binding epitopes recognised by the human monoclonal antibodies formed at different locations in the secretory pathway during S protein transport, but a conformational change in the S1 protein subunit at the Golgi complex formed antibody binding epitopes that are recognised by virus neutralizing antibodies.

## Introduction

Severe acute respiratory syndrome coronavirus 2 (SARS-CoV-2) belongs to the *Coronaviridae* family of viruses, which includes both established coronaviruses that usually cause mild to moderate respiratory disease in humans (e.g., the human coronavirus 229E) [1], and newer emerging viruses that are associated with higher mortality rates in humans (e.g. SARS-CoV-1 and MERS)[2]. The SARS-CoV-2 is a newer emerging virus that are associated with high mortality rates in humans that was first identified in Wuhan [3, 4]. Since the first published sequence of SARS-CoV-2 that was isolated in Wuhan (WIV04), new sequence variants of SARS-CoV-2 have continuously emerged that have exhibited variations in the virus genome sequences that are expected to change the biological properties of the virus. It is predicted that sequence variations may change the virus transmission and virulence characteristics of the in the newly circulating SARS-CoV-2 strains that may impact on control measures that are used to control virus infection [5–7]. Therefore, an improved understanding of the biology of circulating SARS-CoV-2 variants will better clarify the risks that the new sequence variants pose in the human population.

The SARS-CoV-2 genome contains a single non-segmented positive-sense RNA molecule that associates with the N (nucleocapsid) protein to form the virus nucleocapsid which is packaged within the mature virus particle[8]. The mature virus particle has a spherical morphology and is approximately 60-140 nm in diameter, it is surrounded by a lipid envelope into which the virus encoded membrane (M) and spike (S) proteins are inserted[9, 10]. The S protein exists as a homotrimer and protrudes from the virus envelope as an array of club-like projections, and it is a prominent feature on the surface topology of the virus particles[9]. The S protein allows attachment of the virus particle to the cell by binding to the angiotensin-converting enzyme 2 (ACE2) receptor cell receptor. After cell attachment, the S protein also mediates the fusion of the virus and cell membranes that allows the entry of the virus nucleocapsid into the host cell. The S protein is initially synthesized as a single polypeptide chain precursor (S0) that is subsequently cleaved into the S1 and S2 subunits by cellular proteases, and it is this mature form of the S protein containing the two interacting protein subunits that is incorporated into the virus envelope (reviewed in [11]). The S1 subunit contains the ACE2 receptor binding domain (RBD), while the S2 subunit is anchored to the virus envelope by a transmembrane domain and contains the fusion peptide and heptad repeat regions that mediate the process of membrane fusion. The SARS-CoV-2 S protein exhibits some similarities and some differences to the S protein in the previous SARS-CoV-1, such as differences in the cellular protease activity that is involved in processing of the S protein[12].

Due to the importance of the S protein during the initial stages of the virus replication cycle, the S protein has become a focus in the development of antiviral immunological-based strategies. In this context active immunization (vaccines) and passive immunization using therapeutic antibodies (human monoclonal antibodies (hMAbs)) that target the S protein have been deployed to control SARS-CoV-2 infection in the population. Given the importance of the RBD in the S protein during the initial stages of virus cell attachment, these immunological approaches often target the RBD to prevent SARS-CoV-2 interaction with the host cell [13, 14]. We have previously described a panel of S protein hMAbs that were isolated using the B cells of convalescent COVID-19 patients in Singapore[15]. Several of these antibodies that bound to the RBD (RBD-binders) and blocked engagement of the SARS-CoV-2 with the host cell receptor were evaluated as potential therapeutic interventions in SARS-CoV-2-infected patients[16]. Since these antibodies were isolated from individuals recovering from a natural infection, they represent part of a natural polyclonal response to virus infection. Therefore, in addition to their application in passive immunisation, these antibodies also represent physiologically relevant immunological tools that can be employed in experimental cell systems to better understand processing of the SARS-CoV-2 S protein. We have recently described the further characterisation of a subset of the original antibody panel in SARS-CoV-2-infected cells; to better understand their S protein binding characteristics[17]. These antibodies were functionally characterised based on their capacity to neutralise virus infection and to bind to the SARS-CoV-2 S protein RBD. In our earlier study we showed that in virus-infected cells the RBD-binders exhibited an antibody-specific staining pattern that localised within the Golgi complex of infected cells [17]. We had earlier hypothesised that during the export of the S protein through the secretory pathway these antibodies may recognise a conformational change in the S protein in which the RBD becomes surface accessible thus allowing antibody binding to the exposed epitopes. In this current study we have extended our earlier study to address this earlier hypothesis by using these immunological reagents to examine the export of the S protein through the secretory pathway.

## MATERIALS AND METHODS

### Antibodies and specific reagents

The hMAbs PD5, PD4, sc23 and sc29 were isolated using B cell convalescent patients serum and the production of the REGN-10987 antibody were described previously [17]. The rabbit polyclonal antibody to S protein (polyS) was purchased from Sino Biological, Singapore, and the anti-NTD (E7M5X) and anti-RBD (E2T6M) were purchased from Cell Signalling Technology. The anti-rabbit, anti-mouse, and anti-human IgG conjugated to Alexa 488 and Alexa 555 were purchased from Invitrogen (Thermo Fisher Scientific). The Golgi (RCAS1) and endoplasmic reticulum (PDI) as part of the Organelle Localization IF Antibody Sampler Kit #8653 were purchased from Cell Signalling Technology. The Wheat germ agglutinin conjugated to Alexa Fluor™ 488 (WGA-AL488) was purchased from Thermo Fisher Scientific, and Alexa Fluor^TM^ 488 phalloidin was purchased from Cell Signalling Technology.

### Recombinant SARS-CoV-2 S protein expression

The construction of the S protein expression vectors has been described previously [17]. Briefly, the S gene of SARS-CoV-2 (accession no. MN908947) was synthesized by Twist BioSciences (San Francisco, CA, USA). The codon-optimized full-length S gene (nucleotide residues 1 to 1273) was assembled as previously described [17] and cloned into the recombinant pCAGGS vector to create pCAGGS/S-FL. For S1 subunit and RBD expression, only residues 1 to 685 (S1) and 331 to 524 (RBD) were cloned together with a c-terminal FLAG and c-myc tag respectively. The fragments were cloned into pCAGGS to generate pCAGGS/S1-FLAG and pCAGGS/RBD-cMyc. Bulk preparation of all plasmids was performed using the plasmid Midiprep kit (Qiagen). Vero cells (1 × 10^5^ cells per well) in a 24-well cluster plate were transfected with 1 μg plasmid DNA using the Lipofectamine 3000 system (Invitrogen) following the manufacturer’s instructions. The transfected cells were incubated at 33°C in a humidified chamber with 5% CO2 and at 20 hrs post-transfection the cells were processed further.

### Immunofluorescence microscopy

The cells on 12-mm circular glass coverslips were fixed with 4% paraformaldehyde (Sigma-Aldrich) and washed with PBS. The cells were either nonpermeabilized or permeabilized using 0.1% Triton X-100 in PBS at 4°C for 15 min prior to antibody staining. The cells were stained with the appropriate primary and secondary antibody combinations and mounted on microscope slides using CitiFluor. Surface labelling using Wheat germ agglutinin conjugated to Alexa Fluor™ 488 (WGA-AL488) was performed as follows. The non-permeabilised cells were first antibody labelled after which the cells were permeabilised and stained using WGA-AL488 according to the manufacturer’s instructions. For immunofluorescence microscopy imaging the stained cells were imaged using a Nikon Eclipse 80i Microscope (Nikon Corporation, Tokyo, Japan) with an Etiga 2000R camera (Q Imaging, Teledyne Photometrics, Tucson AZ, USA) attached. The images of immunofluorescence-stained cells were recorded using Q Capture Pro ver. 5.0.1.26 (Q Imaging, Teledyne Photometrics). Imaging for confocal microscopy was performed using a Zeiss 910 confocal microscope (Zeiss, Oberkochen, Germany) with Airyscan 2 processing using the appropriate machine settings. The recorded images were examined and processed using Zen ver. 2.3 software (Zeiss).

### Western blotting

The transfected cells were washed using ice cold PBS (4 °C) and extracted directly into Boiling Mix (1% SDS, 5% mercaptoethanol in 20 mM Tris/HCL, pH 7.5). After heating at 95 °C for 2 min the cell extracts were clarified by centrifugation (13,000×g for 2 min) and the proteins separated by SDS-PAGE and transferred by Western blotting onto nitrocellulose membranes. In all cases the apparent molecular masses were estimated using Kaleidoscope protein standards (Bio Rad, USA). The protein bands were visualized using the ECL protein detection system (Amersham, USA).

## Results and discussion

### 1. The human monoclonal antibodies (hMAb) used in this study

The PD5, PD4, sc23 and sc29 human monoclonal antibodies (hMAb) that were used in this study were originally classified based on their binding characteristics to the purified RBD [15]. The PD5 and sc23 antibodies were representative of the strong and weak RBD-binders respectively, while the PD4 and sc29 antibodies were representative of the antibody class that failed to bind to the RBD. The RBD recognition of these antibodies correlated with their capacity to neutralise virus infection, with the PD5 and sc23 exhibiting virus neutralising activity and the PD4 and sc29 failing to neutralise virus infection. Although these antibodies differed with respect to RBD recognition, they all recognise conformational-specific epitopes in the S protein which has influenced our assay methods used to detect antibody recognition. Detection of their binding to the S protein is incompatible with standard Western blotting techniques since the structural features that give rise to the antibody-binding epitopes in the S protein are disrupted during the sample processing process in e.g. Laemmli sample buffer. The recognition of the purified S protein ectodomain and the RBD by these antibodies was previously demonstrated using ELISA-based assays in which the structural integrity of the S protein is preserved [15, 17]. However, in this current study we have used immunofluorescence (IF) microscopy to examine the antibody-stained cells expressing the S protein as the primary means to analyse the immunoreactivity of the hMAb. This strategy also allowed us to assess the immunoreactivity of these antibodies and to also determine the cellular distribution of the S protein species that these antibodies recognise, thus allowing us to examine antibody binding in an *in situ* cell context that can’t be achieved using purified S protein species. In this current study we examined the antibody binding characteristics of these antibodies by recombinant expression of the S protein using the S protein sequence of the SARS-CoV-2/WIV04 strain [4]. We have also used African green monkey kidney epithelial cells (Vero cells) to express the S protein and examine the binding properties of these antibodies. This is an established permissive cell system used to propagate the SARS-CoV-2 that leads to the production of infectious SARS-CoV-2 particles. In this context important post-translational modification of the recombinant expressed SARS-CoV-2 S protein also occurs in these cells, such as Furin cleavage and N-linked glycosylation. Examining the recombinant expressed S protein in Vero cells also enables a direct comparison to be made with our earlier study that used these antibodies to examine SARS CoV-2-infected Vero cells [17]. SARS-CoV-2-infected Vero cells were co-stained with each of the hMAb and the PolyS antibody which confirmed the staining patterns seen previously (SFig. 1). The PolyS antibody is a commercially available immunological reagent that recognises the S2 protein subunit, and it is compatible for use in both imaging and immunoblotting analyses. The PD5 and sc23 antibodies exhibited a distinct punctate staining pattern in these cells, which contrasted with the more generalised and diffuse PD4 and sc29 antibody staining pattern that was similar to the PolyS antibody staining pattern in these cells.

**Figure 1.**
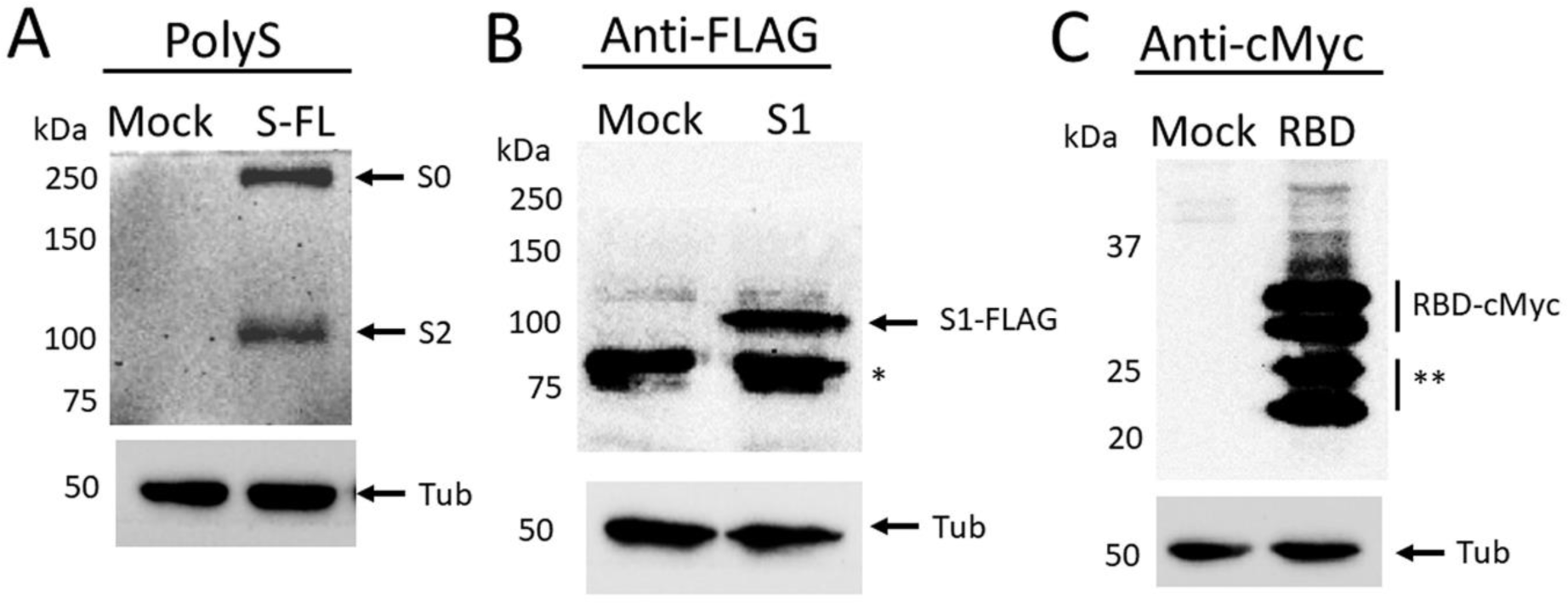
Recombinant expression of the S protein in Vero cells. Cell lysates were prepared from cells transfected with **(A)** pCAGGS/S-FL (S-FL) **(B)** pCAGGS/S1-FLAG (S1) and **(C)** pCAGGS/RBD-cMyc (RBD) and the lysates immunoblotted with the polyS, anti-FLAG and anti-cMyc antibodies respectively and as indicated. In each case the antibodies were also immunoblotted against lysate prepared from mock-transfected cells (Mock). Protein species corresponding in size to the un-cleaved S protein precursor (S0), S2 protein subunit, S1-FLAG protein and RBD-cMyc protein are indicated. Tubulin is the loading control and * is a non-specific protein that appears when immunoblotting with anti-FLAG and ** are the smaller RBD-cMyc species detected in pCAGGS/RBD-cMyc-transfected cells.

### 2. Determining the S protein subunit binding specificity of the hMAb

The full-length SARS-CoV-2 S protein sequence (S-FL), the S1 protein subunit sequence tagged with the FLAG sequence (S1-FLAG) and the RBD sequence tagged with cMyc sequence (RBD-cMyc) were inserted into the plasmid pCAGGS to generate pCAGGS/S-FL, pCAGGS/S1-FLAG and pCAGGS/RBD-cMyc respectively. Cell lysates prepared from cells expressing the S-FL and immunoblotted with PolyS antibody revealed protein species of the expected size for both the S0 precursor and the S2 protein subunit (Fig. 1A). Cells lysates prepared from cells expressing the S1-FLAG and RBD-cMyc were immunoblotted with anti-FLAG and anti-cMyc respectively. An approximate 100 kDa S protein species of the expected size for S1-FLAG protein was detected in cell lysates prepared from pCAGGS/S1-FLAG-transfected cells and immunoblotted with anti-FLAG (Fig. 1B). In cell lysates prepared from pCAGGS/RBD-cMyc-transfected cells and immunoblotted with anti-cMyc we detected four closely space RBD-cMyc protein species at approximately 23/26kDa and 33/35 kDa, the latter being within the size range for the RBD (Fig. 1C). The reason for the appearance of the smaller 23/26kDa RBD doublet species is currently unclear. However, the respective S protein species were not detected in cell lysates prepared from the mock-transfected cells after immunoblotting with the three antibodies, and collectively the immunoblotting analysis confirmed the expression of the respective S protein species in the transfected Vero cells.

Cells were either mock-transfected or transfected with pCAGGS/S-FL and labelled using the PD5, PD4, sc23, sc29, and PolyS antibodies and the stained cells examined using IF microscopy. While no staining was observed in the mock-transfected cells, specific staining was observed in the pCAGGS/S-FL transfected cells stained with all antibodies (SFig. 2). A closer examination of these cell staining pattern indicated that the PD5 and sc23 stained cells again exhibited a prominent punctate staining pattern that contrasted with the more diffuse staining observed in the PD4 and sc29-stained cells (Fig. 2A). In this study we also used the therapeutic antibody REGN-10987 and PolyS as control antibodies to compare with the staining patterns exhibited by the hMAb. The REGN-10987 antibody has accurately defined binding sites at the RBD in the SARS-CoV-2 S protein and blocks virus cell attachment [18, 19], and this was used as a control antibody in our previous study on imaging of SARS-CoV-2-infected cells [17]. In pCAGGS/S-FL transfected cells stained with REGN-10987 we observed a similar staining pattern to that observed in PD5 and sc23-stained cells. Interestingly, the antibody staining pattern exhibited by PD5 and REGN-10987 was also observed using other antibodies in the hMAb panel that recognise the RBD (Tan and Sugrue, unpublished observations). The PolyS-stained cells exhibited a more diffuse staining pattern that was similar to staining pattern exhibited in the PD4 and sc29-stained cells. Although the PD4 and sc29 antibodies exhibited a generalised and diffuse staining pattern across the cell that was distinct from the prominent localised PD5 staining pattern; however, lower levels of an additional more localised staining pattern that was similar to that observed for PD5 was also noted in the PD4 and sc29-stained. This suggested a small degree of overlap of the PD4 and PD5 staining in this region of the cell and further suggested that the S protein species recognised by PD5 and sc23 may also contain the PD4 and sc29 epitopes. This was consistent with our previous imaging analysis made in virus-infected cells when the different staining patterns by these different antibodies were compared[17]. In addition, in this previous study we described Singapore SARS-CoV-2 isolates that exhibited some minor differences in the amino acid sequence of the S protein when compared with that of the S protein sequence of the SARS-CoV-2/WIV04 strain. These sequence difference in the S protein were centred around the location of the furin cleavage site. Although the work described in this study used the S protein sequence of the SARS-CoV-2/WIV04 strain, these respective antibody staining patterns are also observed in cells expressing the recombinant S protein of these Singapore virus isolates (Wu, Tan et al, unpublished observations), suggesting that these staining patterns described above were not virus strain-specific.

**Figure 2.**
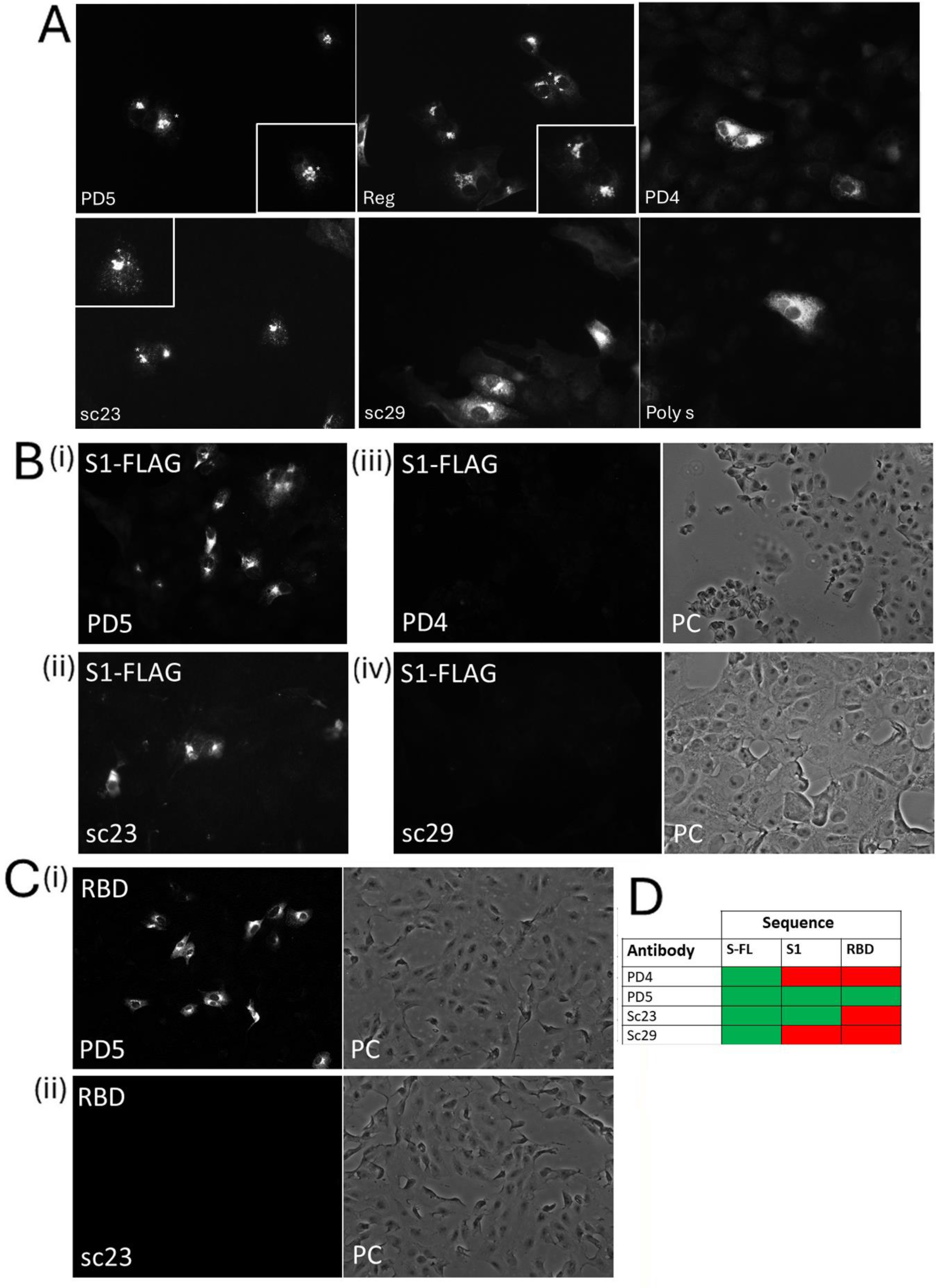
Immunoreactivity of the hMAb panel to the S1 subunit and RBD. **(A)** Cells were transfected with pCAGGS/S-FL and stained using PD5, PD4, REGN-10987 (Reg), sc23, sc29 and polyS as indicated, and then imaged using immunofluorescence (IF) microscopy (objective ×20 magnification). Insets are enlarged images of the individual representative antibody-stained cells showing the distinctive staining patterns. **(B)** Cells were transfected with pCAGGS/S1-FLAG (S1-FLAG) and stained with (i) PD5, (ii) sc23, (iii) PD4, and (iv) sc29 and imaged by IF microscopy and phase contrast microscopy (PC) (objective ×20 magnification) as indicated. **(C)** Cells were transfected with pCAGGS/RBD-myc (RBD) and stained with (i) PD5 and (ii) sc23 and imaged by IF microscopy and phase contrast microscopy (PC) (objective ×20 magnification). **(D)** is a table summarising the immunoreactivity of PD5, sc23, PD4 and sc29 against the entire S protein (S-FL), the S1 subunit (S1) and the receptor binding domain (RBD). Immunoreactive (green box) and non-immunoreactive (red box) are indicated.

The immunoreactivity of the hMAb in cells expressing either the entire S1 domain or the RBD was examined to better understand the S protein recognition by the hMAb. Cells that were transfected with pCAGGS/S1-FLAG and stained using either of the four antibodies and examined using IF microscopy. In this assay antibody staining was noted in cells labelled with PD5 (Fig. 2B(i)) and sc23(Fig. 2B(ii)), confirming that these antibodies recognised a site within the S1 domain. In cells labelled with PD4 (Fig. 2B(iii)) or sc29 (Fig. 2B(iv)) we failed to detect antibody staining, although phase contrast microscopy confirmed the presence of cells in the field of view. On this basis and by default, we therefore concluded that the PD4 and sc29 antibodies bind to the S protein at sites located within the S2 domain rather than the S1 domain. This is consistent with our previous study that demonstrated high binding affinity of the PD4 and sc29 antibodies to the S protein, but that they did not bind to the purified RBD and nor did they exhibit virus neutralising activity [17].

Since only the PD5 and sc23 recognised the S1 subunit, pCAGGS/RBD-cMyc-transfected cells were labelled with PD5 (Fig. 2C(i)) and sc23(Fig. 2C(ii)) and examined using IF microscopy. In this analysis, antibody staining was only observed in the pCAGGS/RBD-cMyc transfected cells labelled with PD5, and we failed to detect any significant levels of sc23 staining in these cells. Collectively these data established that while the PD5 and sc23 recognise the S1 subunit, only the PD5 binds to a region of the S1 that contains the RBD (Fig. 2D). Based on the staining intensity exhibited by the transfected cells we concluded that both sc23 and PD5 exhibited similar binding properties to the S1 domain, which is consistent with sc23 and PD5 exhibiting similar binding affinities for the S protein in our previous study [17]. However, in this current study we failed to detect sc23-staining in the RBD-expressing cells, suggesting that sc23 binds to other regions of the S1 protein subunit that are distinct from the RBD. This is consistent with our previous study which indicated significantly reduced binding affinity of sc23 for the purified RBD when compared with PD5. However, in binding competition assays prior binding of the sc23 to the S protein ectodomain was able to impair PD5 binding, and in addition sc23 also exhibited virus neutralisation activity [17]. Taken together these earlier data and the current analysis suggest that sc23 binds close to the RBD region in the S1 protein subunit, but the efficient binding of sc23 to the S protein may require the interaction of other regions of the S1 protein subunit that are adjacent to the RBD region. These data also suggest that human monoclonal antibodies that bind to the S1 protein subunit at sites other than the RBD may still be able to block the engagement of the S protein with its cell receptor.

### 3. The PD5 antibody recognises a specific conformational change in the S1 protein subunit at the location of the RBD

In cells expressing the recombinant S protein we employed two established commercial antibodies with which to compare with the PD5 and sc23 antibodies, these recognise the N-terminal domain (anti-NTD) and the RBD (anti-RBD) in the S1 protein subunit. The anti-NTD and anti-RBD antibodies have been previously used by others to examine and confirm S protein expression using different experimental systems [20–22]. The anti-NTD and anti-RBD antibodies were produced by using synthetic peptides from sequences in the region of the S protein at Proline25 (in the S1 subunit N-terminal domain) and Serine459 (in and the RBD). It is presumed that anti-NTD and anti-RBD recognise linear epitopes within the S protein, and they would therefore be expected to detect the S1 protein subunit and the RBD respectively without conformational constraints in the S protein structure. These antibodies would therefore be expected to detect the total distribution of the S protein (S1 subunit and RBD) in cells expressing the S protein at all stages during its transport through the secretory pathway. We presumed that if PD5 binding to the S protein was due to the exposure of the RBD during tracking of the S protein through the secretory pathway, then in cells expressing the full-length S protein (S-FL) the anti-RBD would exhibit a similar staining pattern to the PD5 antibody.

Mock-transfected and pCAGGS/S-FL cells were labelled with anti-NTD and anti-RBD and examined using IF microscopy and phase contrast microscopy (Fig. 3A). We only observed antibody staining with these reagents in the pCAGGS/S-FL-transfected cells, confirming their specificity with regards to S protein recognition. To confirm recognition of the respective domains (i.e. NTD and RBD) by the anti-NTD and anti-RBD antibodies, cells that were transfected with pCAGGS/S-FL, pCAGGS/S1-FLAG and pCAGGS/RBD-cMyc and labelled with anti-NTD and anti-RBD were examined by IF microscopy. In this analysis, we also used the PD5 and REGN-10987 antibodies as appropriate controls with which to compare the anti-NTD and anti-RBD staining patterns. All antibodies exhibited staining in the pCAGGS/S-FL transfected cells indicating that all antibodies recognised the full-length S protein (Fig. 3B). However, while the cells stained with PD5 and REGN-10987 exhibited a similar punctate straining pattern, a more widespread and diffuse antibody staining pattern in the cells labelled with either the anti-NTD and anti-RBD was noted and was the expected staining pattern for the general detection of the S protein throughout the cell by these antibodies.

**Figure 3.**
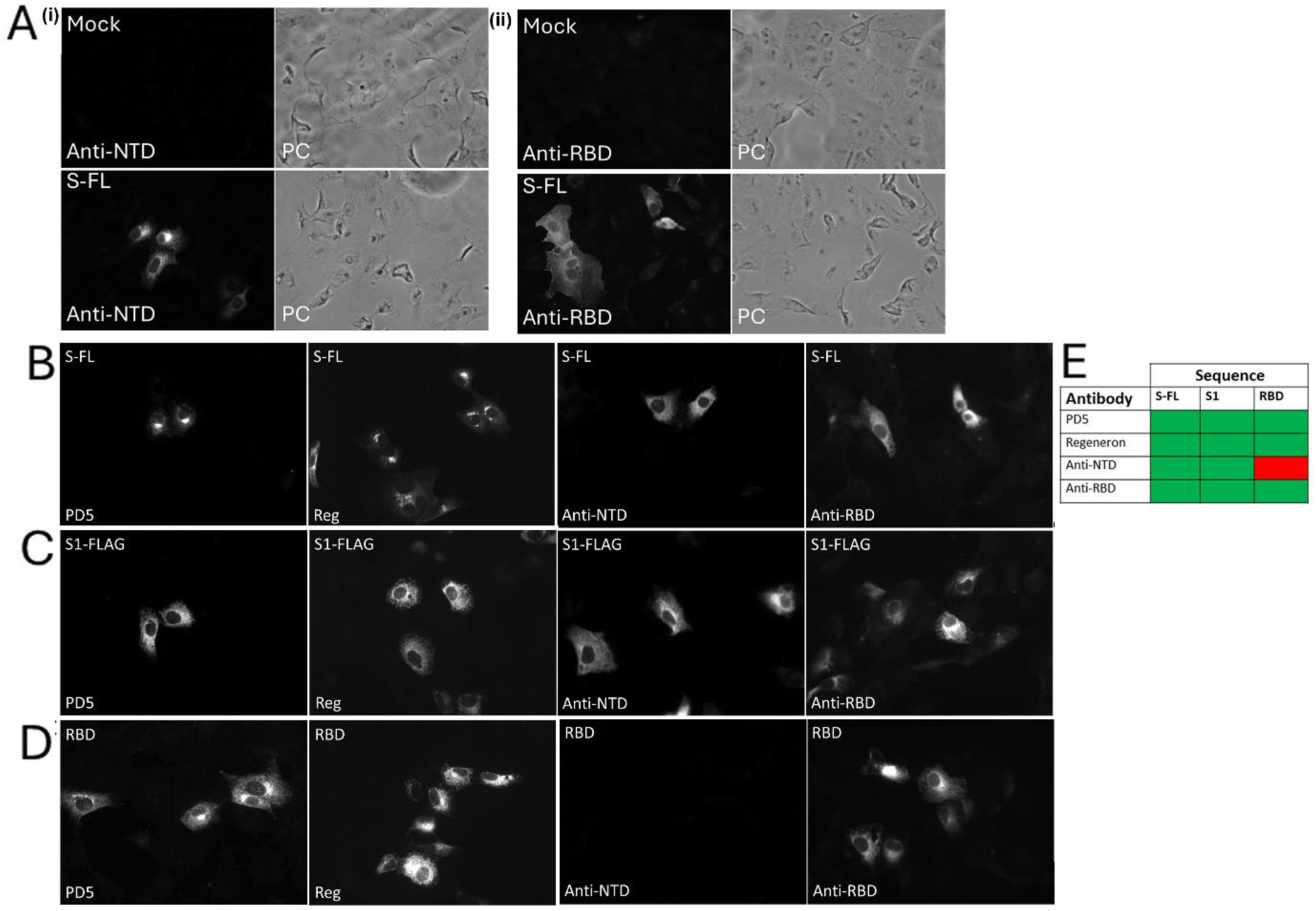
Immunoreactivity of anti-NTD and anti-RBD antibodies. **(A)** Cells were mock-transfected (Mock) and transfected with pCAGGS/S-FL (S-FL) and stained using (i) anti-NTD and (ii) anti-RBD. The cells were imaged using immunofluorescence (IF) microscopy and phase contrast microscopy (PC) (objective ×40 magnification). Cells were transfected with **(B)** pCAGGS/S-FL (S-FL), **(C)** pCAGGS/S1-FLAG (S1-FLAG) and **(D)** pCAGGS/RBD-myc (RBD) and stained with PD5, REGN-10987 (Reg), anti-NTD and anti-RBD as indicated. The cells were imaged by IF microscopy (objective ×40 magnification). **(E)** is a table summarising the immunoreactivity of PD5, Reg, anti-NTD and anti-RBD against the entire S protein (S-FL), the S1 subunit (S1) and the receptor binding domain (RBD). Immunoreactive (green box) and non-immunoreactive (red box) are indicated.

In the pCAGGS/S1-FLAG transfected cells the punctate staining of PD5 and REGN-10987 that was seen in the cells expressing the full-length S protein was absent, and all four antibodies exhibited a similar diffuse antibody staining pattern when examined using IF microscopy (Fig. 3C). This suggested that the prominent punctate staining pattern exhibited by PD5 and REGN-10987 in pCAGGS/S-FL transfected cells required the tethering of the S protein to cell membranes via the membrane anchor in the S2 subunit. In this context, we have also observed a similar diffuse PD5 staining pattern in cells that express the complete S protein ectodomain (without the membrane anchor) (Wu, Tan et al, published observations). This highlights the importance of the S protein transmembrane domain in defining the punctate staining pattern in the cells stained with PD5 and other RBD-binders. However, these data also indicated that membrane association of the S protein was not required for its recognition by PD5 and the other RBD-binders, suggesting that the epitope forms independently of the membrane association of the S protein. Nevertheless, these data confirmed the recognition of the S1 protein subunit by anti-NTD and anti-RBD antibodies. In the pCAGGS/RBD-cMyc transfected cells we observed a diffuse general staining pattern in cells labelled with PD5, REGN-10987 and anti-RBD. As expected, we failed to detect anti-NTD antibody staining in the RBD-cMyc expressing cells (Fig. 3D), which is consistent with the anti-NTD binding to the S protein NTD that is at a location within the S1 subunit that is outside of the RBD.

Collectively these data confirmed that while the anti-NTD and anti-RBD bound to the full-length S protein, anti-NTD and anti-RBD antibodies recognised the NTD in the S1 subunit and the RBD respectively (Fig. 3E). The generalised staining distribution exhibited by anti-NTD and anti-RBD in the pCAGGS/S-FL transfected cells provided evidence that these antibodies were able to detect the presence of the S1 subunit and the RBD at all stages of the transport of the S protein through the secretory pathways. This analysis also confirmed that anti-NTD and anti-RBD would be suitable immunological reagents with which to compare with the specific staining patterns exhibited by the different hMAb in cells expressing the full-length S protein.

Cells were transfected with pCAGGS/S-FL and then co-stained with PD5 and either anti-NTD or anti-RBD (Fig 4A) and co-stained with sc23 and either anti-NTD or anti-RBD (Fig 4B) and imaged using IF microscopy. This confirmed that the prominent PD5 and sc23 staining patterns contrasted with the more diffuse anti-NTD and anti-RBD staining patterns.

**Figure 4.**
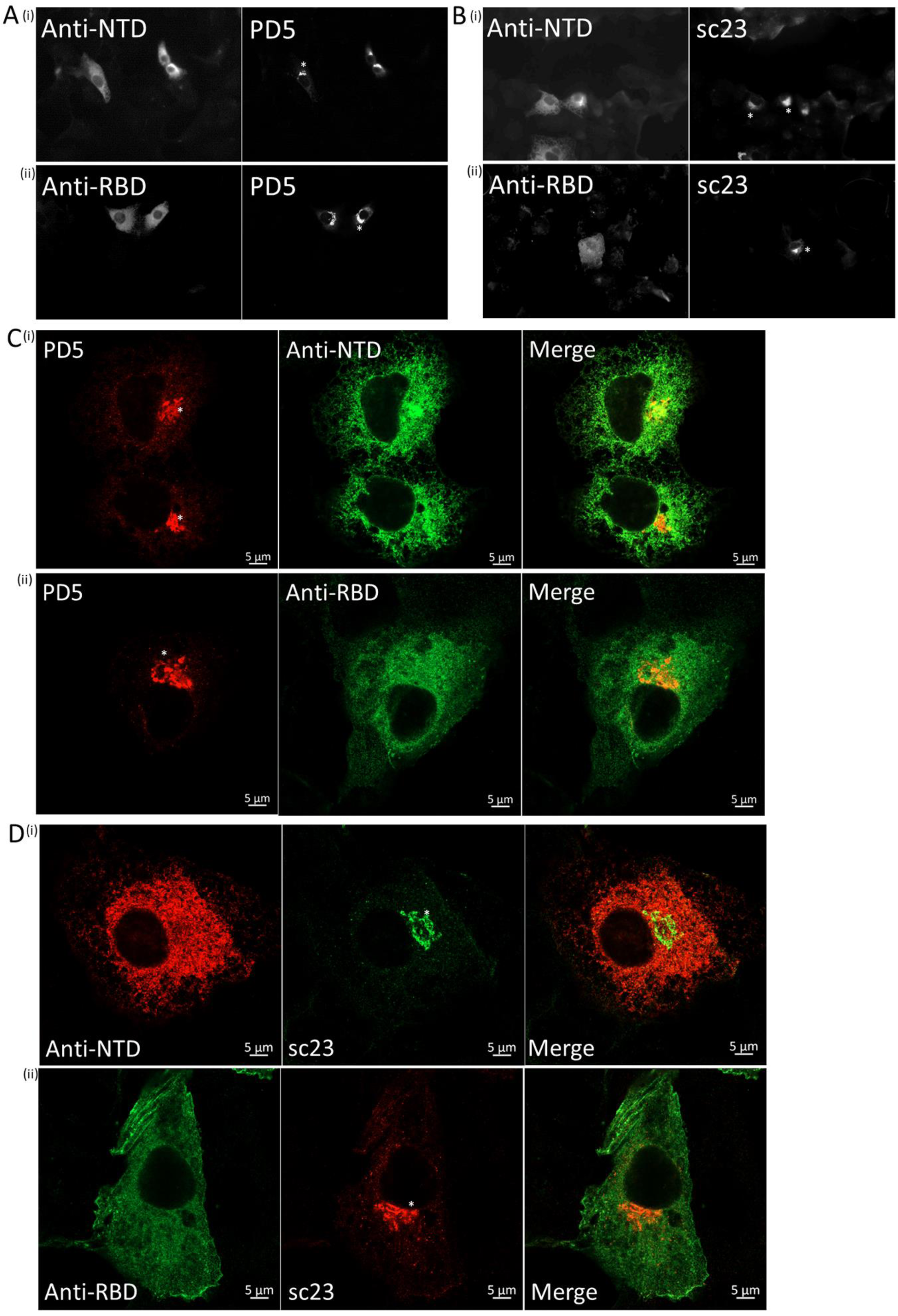
The distribution of the PD5, sc23 and the NTD and RBD staining in cells expressing the S protein. Cells were transfected with pCAGGS/S-FL and co-stained using **(A)** PD5 and (i) anti-NTD and (ii) anti-RBD and **(B)** sc23 and (i) anti-NTD and (ii) anti-RBD and imaged using immunofluorescence (IF) microscopy (objective ×40 magnification). Representative cells that were **(C)** co-stained with PD5 and (i) anti-NTD and (ii) anti-RBD and **(D)** co-stained with sc23 and (i) anti-NTD and (ii) anti-RBD were examined using confocal microscopy. A series of images from the stained cell was obtained in the Z-plane and individual optical slices from the cell interior are shown in each case. In **(C)** and **(D)** individual image channels and the merge image channel are shown. The punctate PD5 and sc23 staining pattern is highlighted (*).

We used confocal microscopy to provide more detailed imaging of the respective antibody staining patterns in individual representative cells co-stained with PD5 and anti-NTD (Fig. 4C(i)) or anti-RBD (Fig. 4C(ii)) and co-stained with sc23 and anti-NTD (Fig. 4D(i)) or sc23 and anti-RBD (Fig. 4D(ii)). In each antibody staining combination we recorded a series of images from individual representative co-stained cells in the Z-plane, and we extracted single images from the image series at an optical plane that allowed us to compare the cytoplasmic staining patterns of the different antibodies. This analysis highlighted the prominent localised punctate PD5 and sc23 staining patterns in these cells, which contrasted with the more generalised and diffuse prominent anti-NTD and anti-RBD staining patterns.

We also performed a detailed analysis of the antibody staining in in the representative co-stained cells by examining the pixel distribution in representative whole cells following PD5 and anti-NTD (Fig. 5A), PD5 and anti-RBD (Fig. 5B) and sc23 and anti-NTD (Fig. 5C) and sc23 and anti-RBD (Fig. 5D) co-staining. The levels of colocalization between the PD5 and the sc23 was compared with either and anti-NTD (Fig. 5A(iii) and C(iii)) and anti-RBD (Fig. 5B(iii) and D(iii)) staining was determined by measuring the Pearson’s correlation coefficient. In all staining combinations within the whole cell, low values for the Pearson’s correlation coefficient (R= between −0.19 and +0.10) were recorded. This was consistent with low levels of general colocalization between the PD5 and sc23 antibody staining and that of either anti-NTD or anti-RBD in these cells. A more detailed analysis was performed in each antibody combination using the correlation coefficient (CC) and weighted correlation coefficient (WCC) to compare the pixel distributions of the anti-NTD and anti-RBD staining with the pixel distribution due to the PD5 and sc23 staining and vice versa. As expected this showed low CC and WCC values when the anti-NTD (CC=0.06 to 0.22; WCC=0.10 to 0.31) and anti-RBD (CC=0.07; WCC=0.08 to 0.13) antibody staining were compared with the PD5 and sc23 antibody staining. This indicated that the PD5 and sc23 staining was largely absent in all the areas of the cell that were stained with either anti-NTD and anti-RBD. However, higher CC and WCC values were recorded when the pixel distribution of the PD5 and sc23 staining were compared with that for the anti-NTD (CC=0.86 to 0.99; WCC=0.91 to 0.99) and anti-RBD (CC=0.99; WCC=0.99). This confirmed that while there was little overall colocalization in the cell between the PD5 and sc23 with the anti-NTD and anti-RBD staining, higher levels of colocalization between the localised punctate PD5 and sc23 staining pattern with the anti-NTD and anti-RBD staining was indicated. Collectively these data provided evidence that the S protein RBD was freely accessible to the antibody staining with anti-RBD, which indicated that the punctate PD5 staining was unlikely to be due to the increased accessibility of the RBD for PD5 antibody binding as we had hypothesised in our earlier study [17]. These data suggested that PD5 was able to detect and bind to a specific conformational change in the structure of the S1 subunit in the region of the RBD. Furthermore, the similar staining patterns exhibited by PD5 and REGN-10987 and other RBD-binders suggests these antibodies also recognise the conformational change in the S1 subunit. However, a similar punctate staining pattern was also observed with the sc23 that does not efficiently bind to the RBD in our cell-based assay suggesting that this conformational change may also involve other regions of the S1 protein subunit that are outside of the RBD.

**Figure 5.**
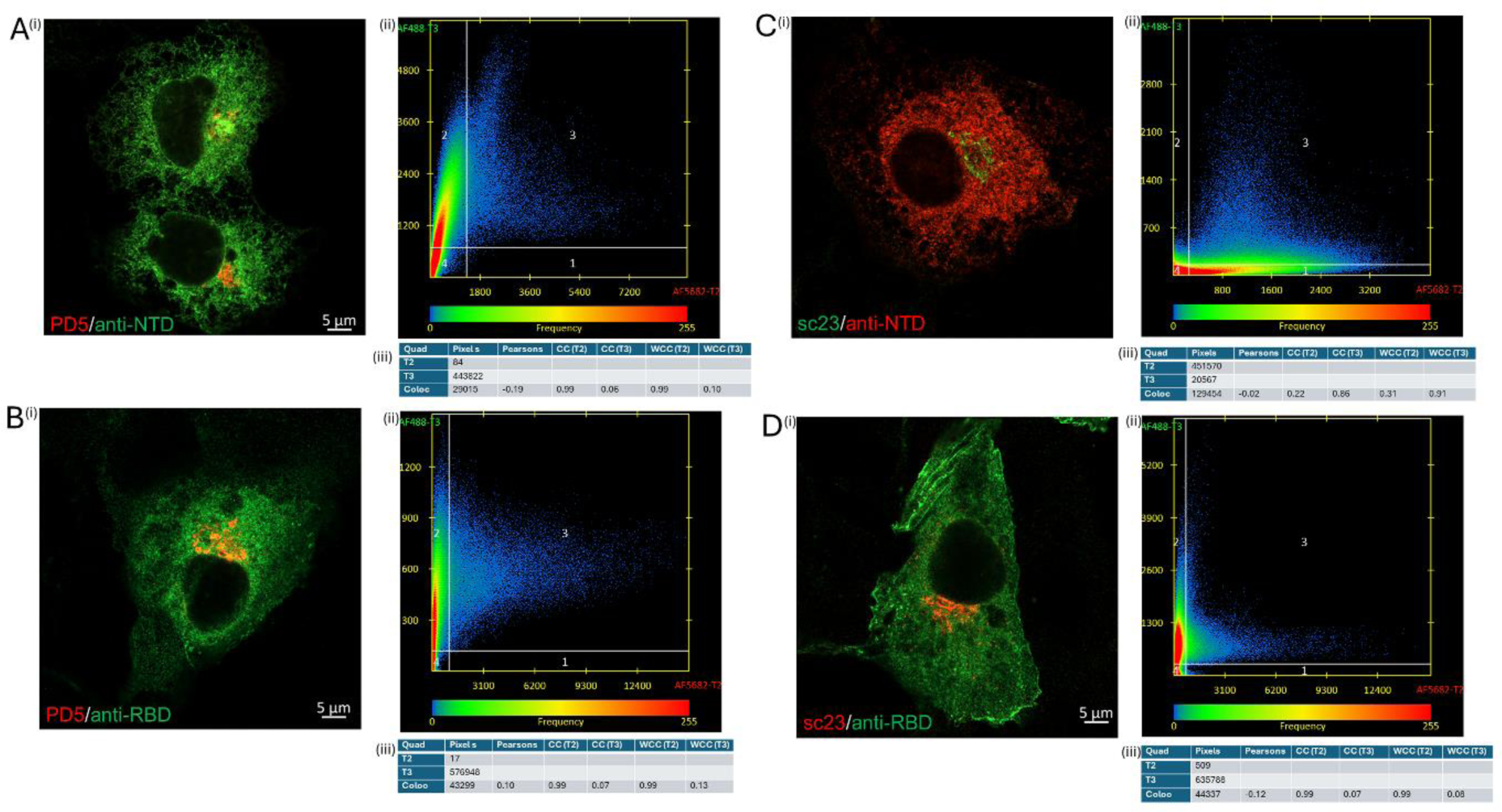
Pixel analysis in the cells co-stained with the NTD and RBD and PD5 and sc23. The pixel distribution in representative cells co-stained with PD5 and **(A)** anti-NTD and **(B)** anti-RBD and sc23 and **(C)** anti-NTD and **(D)** anti-RBD and examined using confocal microscopy were analysed. In each case (i) is the cell examined (merged image only) and (ii) is the resulting scatter plot from that image. (iii) is the table that summarizes the pixel analysis in each antibody combination. The number of pixels only in T2 and T3 are indicated, and (coloc) are the number of pixels used to perform the colocalization analysis. Also shown are the Pearson’s correlation coefficient (Pearsons) for the total number of pixels sampled and the correlation coefficient (CC) and weighted correlation coefficient (WCC) for T2 with respect to T3 (T2) and for T3 with respect to T2 (T3).

Our data suggested that the PD4 antibody recognised a conformational specific epitope that is in the S2 subunit, and we therefore compared the PD4 antibody staining pattern with that of the anti-NTD and anti-RBD staining patterns. The pCAGGS/S-FL transfected cells were co-stained with PD4 and either anti-NTD (Fig 6A (i)) or anti-RBD (Fig 6A(ii))) and imaged using IF microscopy. This revealed the more diffuse and less-localised PD4 staining pattern (compared with PD5 and sc23) and the diffuse anti-NTD and anti-RBD staining patterns. Detailed imaging of the respective antibody staining patterns using confocal microscopy of individual representative PD4 and anti-NTD (Fig. 6B(i)) and PD4 and anti-RBD (Fig. 6B(ii)) co-stained cells highlighted the respective antibody staining patterns of these antibodies. The analysis of the pixel distribution in the PD4 and anti-NTD (Fig. 6C) and anti-RBD (Fig. 6D) staining combinations showed higher CC and WCC values for the anti-NTD (CC=0.54; WCC=0.77) (Fig. 6C(iii)) and anti-RBD (CC=0.3; WCC= 0.34) (Fig. 6D(iii)) staining compared with results of the corresponding analysis with the PD5 and sc23 antibodies described above. When the PD4 staining was compared with that of the anti-NTD and anti-RBD, still higher values for the CC (CC=0.97; WCC=0.98) and the WCC (CC=0.97; WCC=0.98) were again recorded. The pixel analysis indicated higher levels of general PD4 staining in the areas of the cell that were also stained with anti-NTD and anti-RBD and was consistent with the diffuse PD4 antibody staining observed in the cells expressing the S protein.

**Figure 6.**
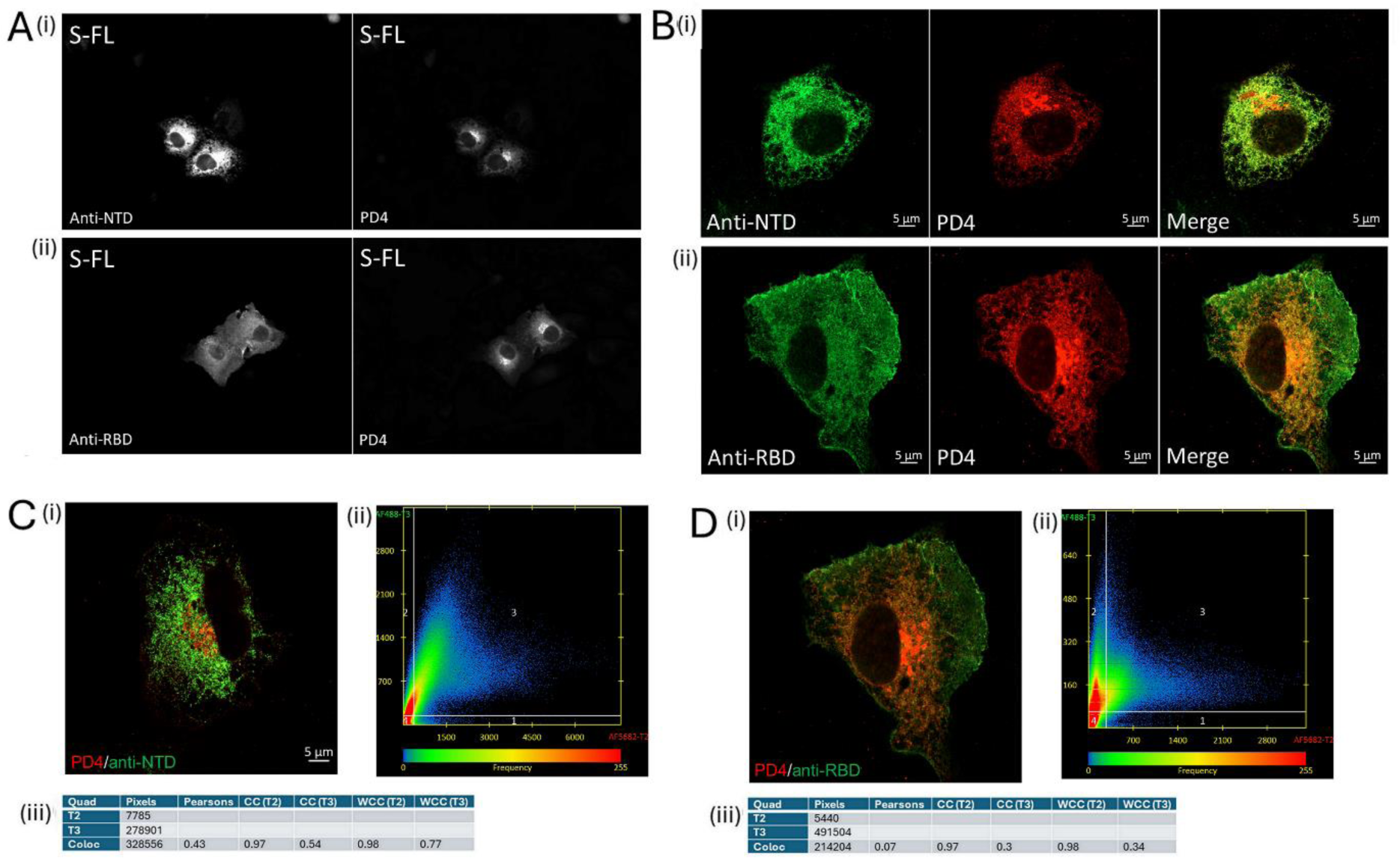
Distribution of the PD4 and NTD and RBD staining in cells expressing the S protein. **(A)** Cells were transfected with pCAGGS/S-FL and co-stained using PD4 and (i) anti-NTD and (ii) anti-RBD and imaged using immunofluorescence (IF) microscopy (objective ×40 magnification). **(B)** Representative cells that were co-stained with PD4 and (i) anti-NTD and PD4 and (ii) anti-RBD were examined using confocal microscopy. A series of images from the stained cell was obtained in the Z-plane and individual optical slices from the cell interior are shown. The individual image channels and the merge image channel are shown. **(C and D)**. The pixel distribution in representative cells co-stained with PD4 and **(C)** anti-NTD and **(D)** anti-RBD were analysed. In each case (i) is the cell examined (merged image only) and (ii) is the resulting scatter plot from that image. (iii) is the table that summarizes the pixel analysis. The number of pixels only in T2 and T3 are indicated, and (coloc) are the number of pixels used to perform the colocalization analysis. Also shown are the Pearson’s correlation coefficient (Pearsons) for the total number of pixels sampled and the correlation coefficient (CC) and weighted correlation coefficient (WCC) for T2 with respect to T3 (T2) and for T3 with respect to T2 (T3).

### 4. The conformational change in the S1 protein subunit that is recognised by PD5 occurs at the Golgi complex

The pCAGGS/S-FL transfected cells expressing the S protein were co-stained with PD5 anti-RCAS1 (which detects the Golgi marker RCAS1) and examined using IF microscopy (Fig 7A). A comparison of the prominent punctate PD5 staining pattern and the anti-RCAS1 staining pattern suggested that these staining patterns coincided i.e. they were at the same location in the cell. Representative PD5 and anti-RCAS1 (Fig. 7A(i)) and sc23 and anti-RCAS1 (Fig. 7B) co-stained cells were examined in greater detailed by using confocal microscopy to examine the respective antibody staining patterns. This analysis provided evidence that the punctate PD5 and sc23 staining localised with the anti-RCAS1 indicating that these were primarily localised at the Golgi complex. Cells expressing the S protein that were also co-stained with PD5 and anti-PDI and examined by IF microscopy (Fig. 7C(i)) and confocal microscopy (Fig 7C(ii))). The anti-PDI detects protein-disulphide isomerase (PDI) which is a protein that is resident in the endoplasmic reticulum (ER) and is an established cellular protein marker that is used for the ER detection. The clearly distinct PD5 and anti-PDI staining patterns demonstrated that the PD5 punctate staining pattern was not associated with the ER. Collectively, these data suggested that the conformational change that leads to the formation of the antibody binding epitopes recognised by the PD5 and sc23 antibodies were not present in the early compartments of the secretory pathway, but these form at the Golgi complex and may accumulate in this cell compartment prior to export to the cell surface. The appearance of the PD5 and sc23 punctate antibody staining pattern at the Golgi complex is also observed in SARS-CoV-2-infected cells [17]. Therefore, the appearance of these staining patterns at the Golgi complex in both virus-infected cells and in cells expressing the recombinant S protein suggests that this localisation was independent of virus infection. This is consistent with several recent studies that have provided evidence that sequence motifs within the SARS-CoV-2S protein are determinants in the trafficking and cellular localisation of the S protein [23–25].

**Figure 7.**
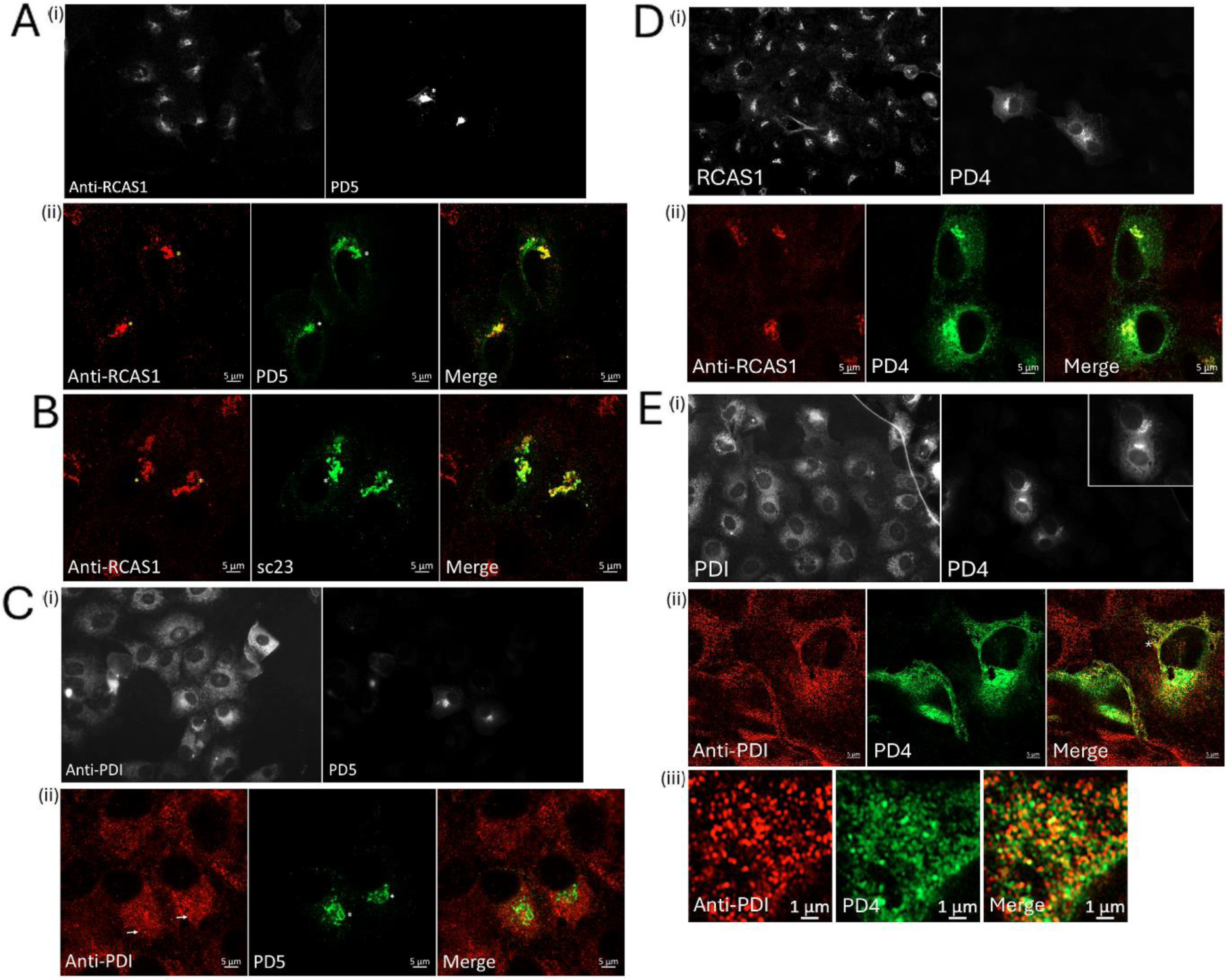
The punctate staining exhibited by PD5 and sc23 localises at the Golgi complex. Cells were transfected with pCAGGS/S-FL and co-stained using **(A)** PD5 and anti-RCAS1 and imaged using (i) immunofluorescence (IF) microscopy (objective ×20 magnification) and (ii) representative PD5 and anti-RCAS1 co-stained cells using confocal microscopy. **(B)** Representative sc23 and anti-RCAS1 co-stained cells were examined using confocal microscopy. The individual image channels and the merge image channel are shown. The punctate PD5 and sc23 staining pattern is highlighted (*). **(C)** Cell were transfected with pCAGGS/S-FL and co-stained using PD5 and anti-PDI and imaged using (i) IF microscopy (objective ×20 magnification) and (ii) representative cells that were co-stained with PD5 and anti-PDI were examined using confocal microscopy. The punctate PD5 staining pattern (*) and the anti-PDI staining (white arrow) are highlighted. **(D)** Cells were transfected with pCAGGS/S-FL and co-stained using PD4 and anti-RCAS1 and imaged using (i) IF microscopy (objective ×20 magnification) and (ii) using confocal microscopy. The individual image channels and the merge image channel are shown. **(E)** Cells were transfected with pCAGGS/S-FL and co-stained using PD4 and anti-PDI and imaged using (i) IF microscopy (objective ×20 magnification) and (ii) by using confocal microscopy. The individual image channels and the merge image channel are shown (iii) is an enlarged image taken from (ii) in the area highlighted (*). In all images recorded using confocal microscopy a series of images from representative stained cell was obtained in the Z-plane and individual optical slices from the cell interior are presented.

We also compared the PD4 antibody staining pattern with that of the anti-RCAS1 (Fig. 7D) and anti-PDI (Fig. 7E) antibodies, to better localise the PD4 staining within the cell. Although the PD4 antibody exhibited a diffuse staining pattern, there was a small degree of overlap with the anti-RCAS1 staining and PD4 staining, suggesting that a small proportion of the S protein species detected by the PD4 antibody was also localised at the Golgi complex. This was consistent with the apparent low level of overlap in the antibody staining patterns exhibited by the PD4 and PD5 antibodies as described above. However, a higher level of co-staining between the diffuse PD4 staining and anti-PDI staining was consistent with the S protein displaying the PD4 epitope in the ER compartment of the cell suggesting that the PD4 antibody binding epitope forms at an earlier compartment of the secretory pathway. Collectively these data provide evidence that the structural changes in the S protein that lead to the formation of the different antibody binding epitopes occur at different stages during the transport of the S protein through the secretory pathway.

### 5. The S protein displaying the PD5 epitope is displayed on microvilli on the surface of cells expressing the recombinant S protein

Although PD5, sc23, PD4 and sc29 display specific cellular staining patterns, we examined if the antibody binding epitopes recognised by these antibodies were also displayed on the surface of cells expressing the S protein. In each case a similar diffuse staining pattern with each of the four antibodies was recorded on the surface of the non-permeabilised pCAGGS/S-FL-transfected Vero cells (Fig. 8A), indicating that these antibody binding epitopes were present on the mature S protein that is trafficked to the cell surface. These staining patterns are distinct from the cytoplasmic staining patterns observed with each of the four antibodies that is described above. These distinctly different surface staining patterns are also seen in non-permeabilised cells SARS-CoV-2-virus infected cells that are stained with each of the four antibodies.

**Figure 8.**
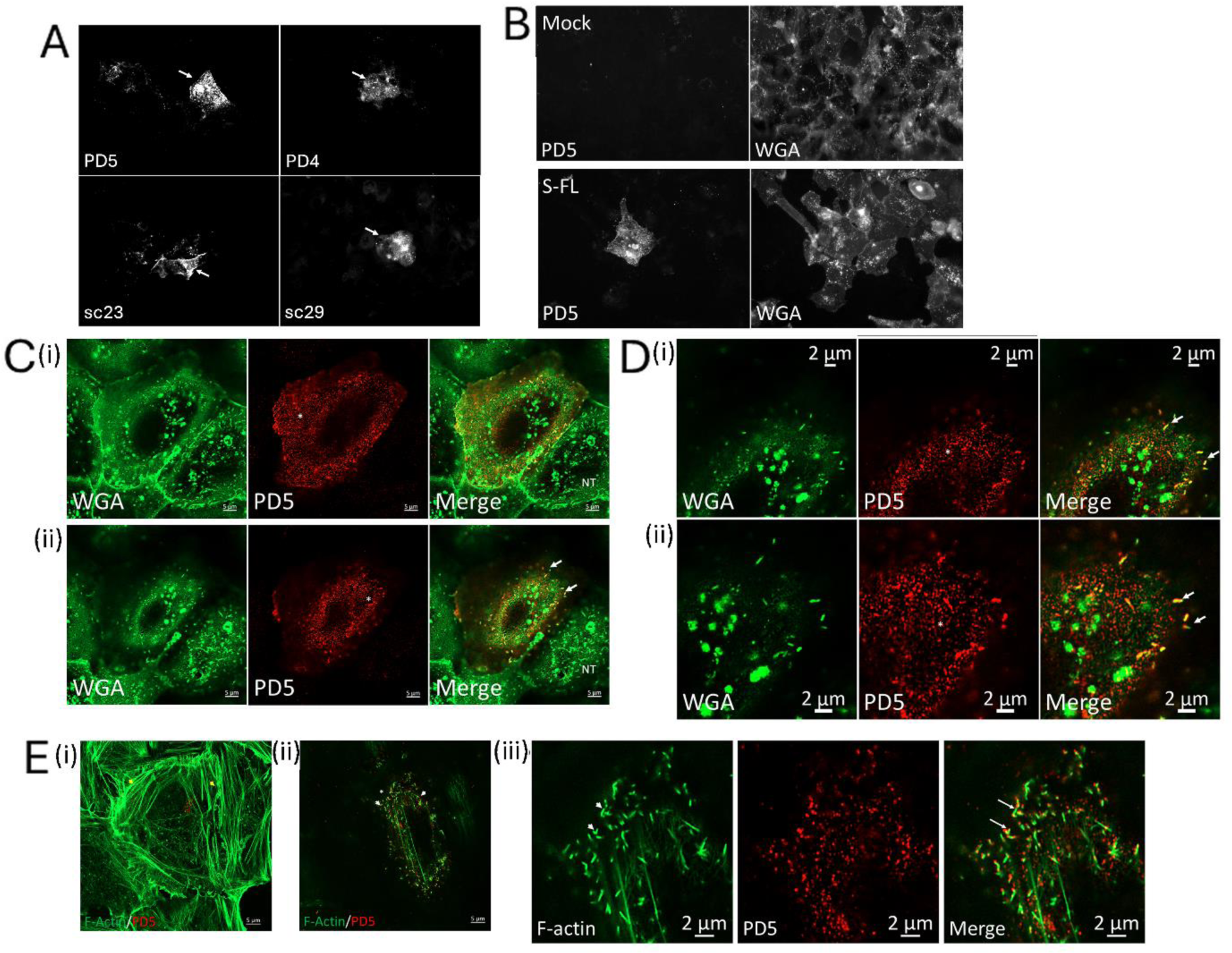
Distribution of the S protein on the surface of pCAGGS/S-FL-transfected cells. **(A)** Cells were transfected with pCAGGS/S-FL and the non-permeabilised cells stained using the PD5, PD4, sc23 and sc29 antibodies and imaged using immunofluorescence (IF) microscopy (objective ×20 magnification). (**B**) Cells were mock-transfected and pCAGGS/S-FL-transfected and the non-permeabilised cells were co-stained with PD5 and wheat germ agglutinin conjugated to Alexa Fluor™ 488 (WGA) and imaged using IF microscopy (objective ×40 magnification). **(C and D)**. Non-permeabilised cells pCAGGS/S-FL-transfected cells were co-stained with PD5 and wheat germ agglutinin conjugated to Alexa Fluor™ 488 (WGA) and examined by confocal microscopy. **(C)** A series of images from the stained cell was obtained in the Z-plane and individual optical slices showing the (i) cell periphery and (ii) cell top are shown. **(D)** are enlarged images showing the cell top. In both **(C)** and **(D)** The individual image channels and the merge image channel are shown and the co-stained filamentous surface projections are highlighted (white arrows). **(E)** pCAGGS/S-FL-transfected cells were co-stained with PD5 and Alexa Fluor^TM^ 488 phalloidin (F-actin) and examined by confocal miscopy. A series of images from the stained cell was obtained in the Z-plane and individual optical slices showing (i) the cell periphery and (ii) the cell top. Only the merged image is shown. (iii) is an enlarged image showing the cell top. The individual image channels and the merge image channel are shown. The presence of the Alexa Fluor^TM^ 488 phalloidin stained filamentous projection (short arrows) and the PD5 staining on these projections (long arrows) are highlighted.

In a final analysis we therefore used the PD5 as a representative antibody in the antibody panel to examine the surface expression of the recombinant S protein on Vero cells. Wheat germ agglutinin conjugated to Alexa Fluor™ 488 (WGA-AL488) is an established cellular probe that is used to image the surface of cells, and we used WGA-AL488 as a probe to visualise the surface topology of the transfected cells. Cells were either mock-transfected or transfected with pCAGGS/S-FL and the non-permeabilised cells labelled with PD5 and WGA-AL488. The WGA-AL488 stained the surface of both the mock-transfected and pCAGGS/S-FL-transfected cells, while PD5 staining was only observed on the pCAGGS/S-FL-transfected cells (Fig. 8B). The PD 5 staining exhibited a diffuse staining pattern over the surface of these cells, which contrasted with the prominent punctate PD5 cellular staining pattern observed in permeabilised cells. These distinct PD5 staining patterns were also observed in PD5-stained SARS-CoV-2-infected cells, suggesting that the correctly folded S protein may accumulate at the Golgi complex prior to its export to the surface of cells expressing the recombinant S protein and on virus-infected cells.

Representative PD5 and WGA-AL488 co-stained cells were examined in more detail using confocal microscopy at a focal plane that allowed imaging of the surface topology of these cells and the distribution of the PD5 surface-staining (Fig. 8C and D). At higher magnification (Fig. 8D(i) and (ii)) we observed filamentous WGA-AL488 surface staining pattern that we interpreted as staining of cell surface projections such as microvilli. Two distinct PD5 staining distributions were noted, one that was present on the smooth areas of the cell surface, and one that was on the WGA-AL488-stained filamentous projections (microvilli). In this context, the S protein displaying the PD5 epitope was displayed on the cell surface of pCAGGS/S-FL-transfected cells in a manner similar to the distribution of virus particles that form on SARS CoV-2-infected cells [17]. We also complimented this assay with an additional analysis by comparing the PD5 surface-staining and Alexa Fluor^TM^ 488 phalloidin staining pattern on pCAGGS/S-FL-transfected cells (Fig. 8E). The Alexa Fluor^TM^ 488 phalloidin stains the cells F-actin network and allows imaging of cell surface projections that are stabilised by short F-action bundles (e.g. in the microvilli). The non-permeabilised cells were first labelled with the PD5 antibody, after which the cells were permeabilised and stained using Alexa Fluor^TM^ 488 phalloidin. We used confocal microscopy to examine representative PD5 and Alexa Fluor^TM^ 488 phalloidin co-stained cells in more detail at a focal plane that allowed imaging of the cell periphery and at the cell top. Imaging of the cell periphery showed the presence of F-actin stress fibres (Fig. 8E(i)), while imaging at the cell top allowed visualisation of the F-actin stabilized cell projections such as the microvilli (Fig. 8E(ii)). A more detailed analysis at the cell top showed the presence of a relatively high level of PD5 staining along the Alexa Fluor^TM^ 488 phalloidin stained microvilli (Fig. 8E(iii)). Therefore, the surface-stained PD5 stained cells that were stained using WGA-AL488 and Alexa Fluor^TM^ 488 phalloidin are consistent and indicated the presence of the S protein on the cell surface microvilli projections. These data demonstrate that although the PD5 epitope forms in the Golgi complex, the S protein displaying this epitope is trafficked to the cell surface and is also trafficked into the microvilli. In this context, high-resolution imaging of SARS-CoV-2-infected Vero cells shows that high levels of virus particles are present on microvilli on the surface of infected Vero, which suggested that microvilli may facilitated localised virus transmission in the cell monolayers [17]. Our current analysis suggests that the S protein is trafficked independently to sites on the cell surface that are associated with SARS-CoV-2 virus particle assembly and virus transmission.

As with other members of the *Coronaviridae* family of viruses, the SARS-CoV-2 particle assembly begins in the early part of the Golgi complex (reviewed in[26]). In this context our data using recombinant expressed S protein is consistent with the transport of the S protein from the Golgi complex into the microvilli during virus infection. Trafficking signals in the c-terminal tail of the coronavirus S protein have been identified [25, 27] and in a final analysis we also examined if the S1-FLAG protein displaying the PD5 antibody binding epitope was also trafficked to the surface of pCAGGS/S1-FLAG transfected cells. Non-permeabilised mock-transfected cells (Fig. 9A), pCAGGS/S-FL-transfected cells (Fig. 9B), and pCAGGS/S1-FLAG transfected cells (Fig. 9C) were co-stained using PD5 and WGA-AL488 and examined by IF microscopy. The WGA-AL488 surface-staining was observed in all three assay conditions, while PD5 staining was only detected on the pCAGGS/S-FL and pCAGGS/S1-FLAG transfected cells. Analysis of the staining patterns on pCAGGS/S-FL-transfected cells showed a diffuse PD5 staining pattern, while a more punctate distinct PD5 staining pattern was observed on the surface of the pCAGGS/S1-FLAG transfected cells. Since the correctly folded S1 protein subunit is expected to be a prerequisite for protein export through the secretory pathway[28, 29], the presence of the S1-FLAG protein on the cell surface indicated that it is trafficked through the secretory pathway and therefore correctly folded and export competent. However, as the S1-FLAG protein does not contain a transmembrane anchor its expression on the cell surface was not expected. Currently the molecular basis for its surface expression is not clear but it may be mediated by the interaction between the S1-FLAG protein and one or more cellular factors on the surface of these cells. We used confocal microscopy to examine PD5 and WGA-AL488 staining on representative co-stained pCAGGS/S1-FLAG transfected cells in more detail, at a focal plane that allowed imaging of the WGA-AL488-stained surface topology and the distribution of the PD5 staining (Fig. 9D). Although relatively high levels of the PD5 staining were observed on the surface of these cells, we failed to detect the presence of significant level of PD5 staining on the WGA-AL488-stained cell surface projections e.g. microvilli. This suggested that while the S1 subunit protein could be trafficked to the surface of these cells independently of the S2 protein subunit, trafficking of the S protein to the microvilli may require the presence of the S2 protein subunit. Although the c-terminus cytoplasmic sequence of the S protein facilitates surface expression of the S protein[23], our data also suggests that structural motifs may be present in the S1 subunit that allow export of the correctly folded S1 protein subunit. These data also suggest that the S1 protein subunit containing the conformational change that is recognised by RBD-binders such as PD5 is exported through the secretory pathway to the cell surface independently of the S2 protein subunit.

**Figure 9.**
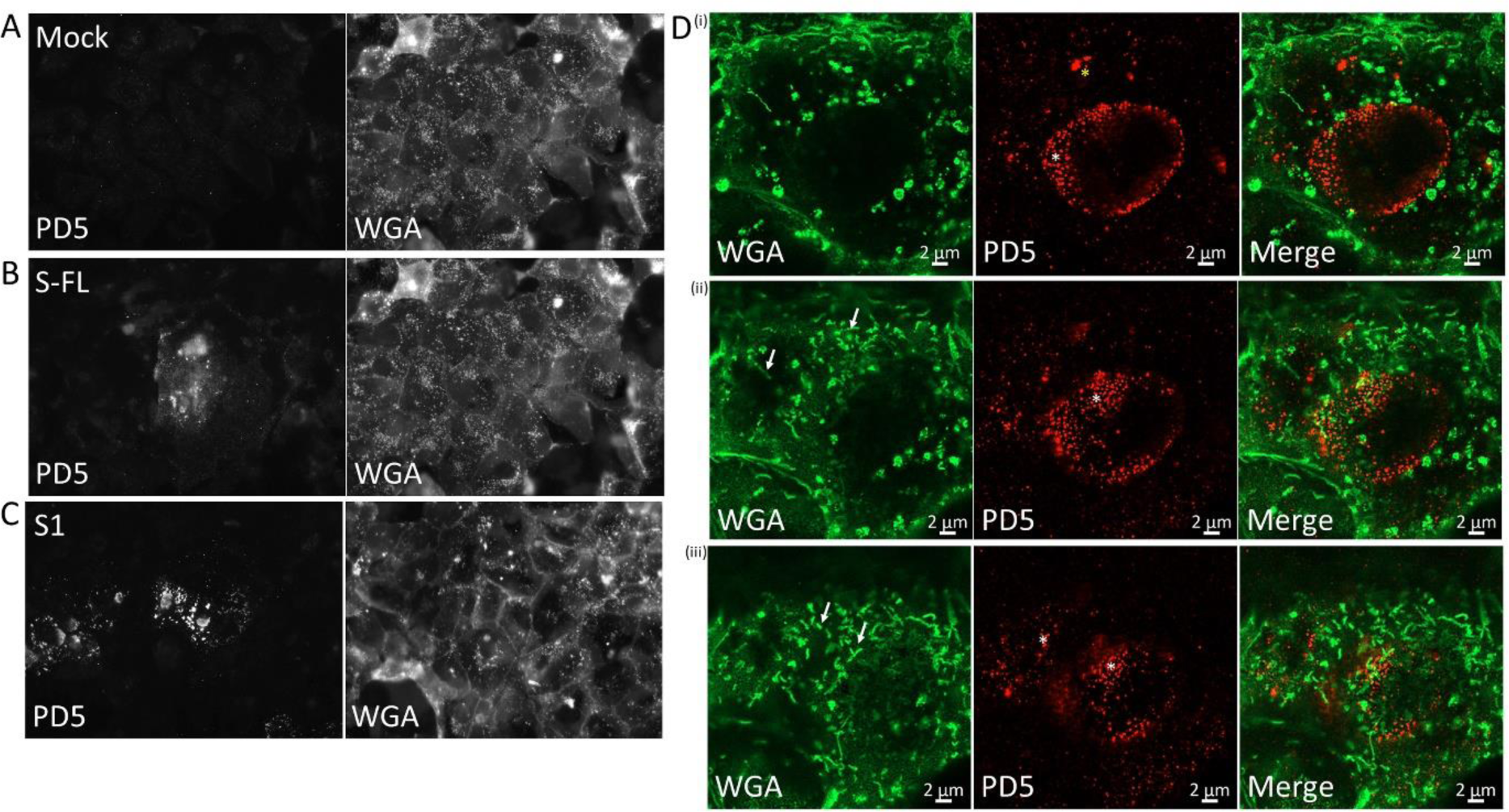
Distribution of the S1 protein subunit on the surface of pCAGGS/S1-transfected cells. Cells were **(A)** mock-transfected, **(B)** transfected with pCAGGS/S-FL (S-FL and **(C)** transfected with pCAGGS/S1 (S1) and the non-permeabilised cells were co-stained with PD5 and wheat germ agglutinin conjugated to Alexa Fluor™ 488 (WGA) and imaged using immunofluorescence (IF) microscopy (objective ×40 magnification). **(D)** Representative non-permeabilised cells transfected with pCAGGS/S1 were co-stained with PD5 and wheat germ agglutinin conjugated to Alexa Fluor™ 488 (WGA) were imaged using confocal microscopy. A series of images from the stained cell was obtained in the Z-plane and individual optical slices at (i) the cell periphery, (ii) close to the cell top, and (iii) at the cell top are shown. The individual image channels and the merge image channel are shown. The WGA-stained filamentous projections on the cell surface (white arrows) and the PD5 staining pattern (*) are highlighted.

## 6. Conclusions

The antibody-specific staining patterns that we observed using the hMAb in this current study were also observed previously in virus-infected cells, suggesting that the processing of the S protein in permissive cells occurs independently of virus infection. This current study demonstrated that these conformational specific human monoclonal antibodies can be used to examine S protein processing *in situ*, and to examine how neutralising epitopes form in the S protein form in virus-infected cells. In our earlier study we had originally hypothesised that during its transport through the secretory pathway, changes in the S protein structure may result in the increased surface exposure of the RBD that allowed the binding of PD5 and other similar RBD-binders[17]. However, in the current study we show that the RBD is accessible to antibody staining throughout the cell, suggesting that the RBD-binders (e.g. PD5 and REGN-10987) recognise a specific 3-dimensional structural conformation in the S1 protein subunit that is formed at the location of the RBD rather than by the recognition of the RBD per se. Antibody binding at these locations in the S protein would be expected to exert a steric hindrance that would prevent its engagement with the cell receptor and block infection. Our data suggested that the structural change in the S1 protein subunit that led to the formation of the antibody binding epitopes that are associated with virus neutralising activity occurred at the Golgi complex, as the S protein was trafficked through the secretory pathway. The processes that underpin this conformational change is currently unclear, but may be associated (directly or indirectly) with biochemical activities in the Golgi complex that lead to the post-translational modification of the S protein e.g. S protein N- and O-linked glycosylation[30–32]. The importance of the structural conformation in the S1 protein subunit that is detected by the RBD-binders to the function of the S protein during virus infection also remains unclear. In addition, PD5 staining was detected on the surface of cells expressing the recombinant S protein, and it is unclear if the conformational change in the S1 protein subunit that was detected by the RBD-binders (e.g. PD5) was required for the efficient trafficking of the S protein from the Golgi complex to the cell surface. If this structure is vital to the functioning of the S protein during virus infection (e.g. by mediating cell attachment) this may have implications for the emergence of antibody escape mutations that may arise in newer circulating virus strains. Mutations in the S protein amino acid sequence that destabilise this structure may lead to the production of a S protein with impaired functionality, while mutations that do not stabilise this structure but that reduce antibody binding activity may be advantageous to the virus and allow immune evasion. If this is the case, then it should be possible to use structure-based analysis of the S protein to predict the sequence of the antibody resistant virus strains that emerge following specific antiviral interventions using therapeutic antibodies and vaccines. In this context the cellular virology approach described in this study should complement the high-resolution structural studies that are currently being employed to examine the interaction of the S protein and therapeutic antibodies. Therefore, future studies will focus on identifying the precise binding sites on the S protein ectodomain for the human monoclonal antibodies described in this study to more accurately define the conformational changes that occur in the S protein in virus-infected cells during virus particle assembly.

## Acknowledgements.

Yanjun Wu is a CN Yang Scholar and was supported by the CN Yang Programme at the Nanyang Technological University. We thank Jie Ling Tan for technical support in the data acquisition and Angeline Pei-Chiew Lim and Steven Ka-Khuen Wong for assistance in antibody preparation. This research received no specific grant from any funding agency in the public, private or not-for-profit commercial sectors.

