## supplementary data for "Evidence that the SARS-CoV-2 S protein undergoes a conformational change at the Golgi Complex that leads to the formation of virus neutralising antibody binding epitopes in the S1 protein subunit"

**Supplementary Figures.**

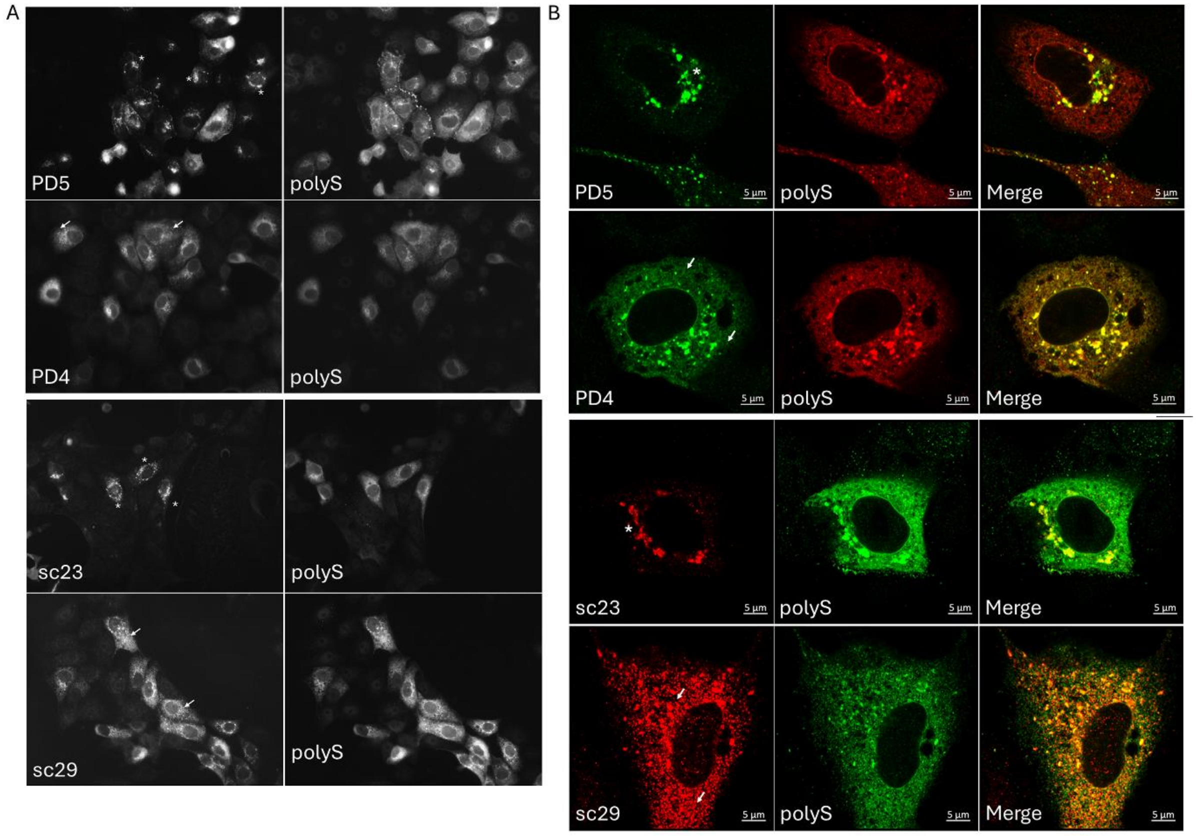


**SFigure. 1 Distribution of the S protein in virus-infected Vero E6 cells stained using the hMAb panel.** Vero E6 cells were seeded onto 12-mm circular glass coverslips and infected with SARS-CoV-2 at the required multiplicity of infection of 0.1 at 37°C as described previously (Chan, et al J Virol, 2022. 96(13): p. e0045522). At 18 hrs post infection the cells co-stained with PolyS and PD5, PD4, sc23 and sc29 as indicated and imaged using **(A)** immunofluorescence microscopy (objective x40 magnification) and **(B)** confocal microscopy (the individual and merged image channels are shown). The punctate PD5 and sc23 staining patterns (*) and the diffuse generalised PD4 and sc29 staining patterns (white arrows) are indicated.

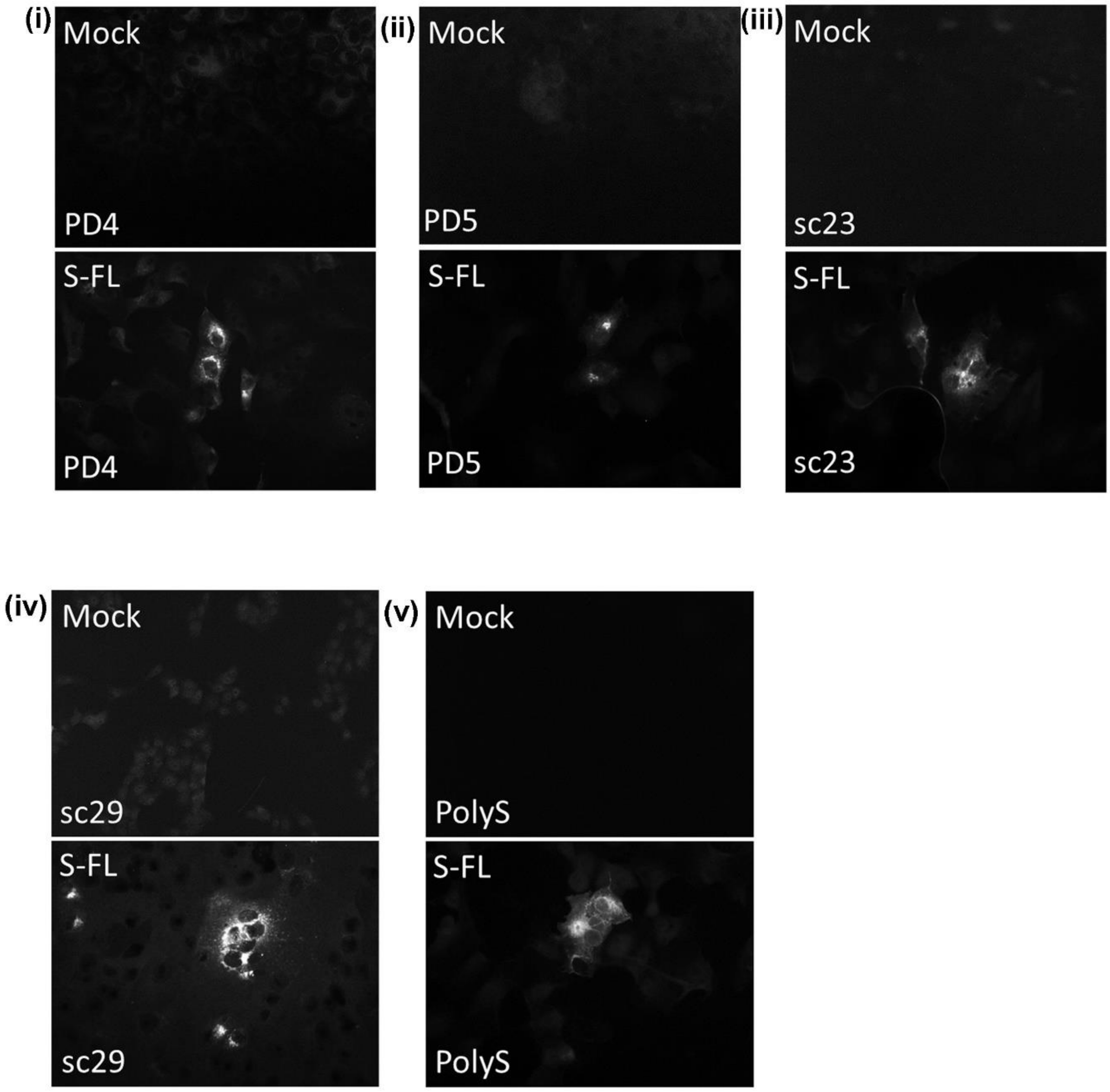


**SFigure 2. Immunoreactivity of the hMAb in c ells expressing the S protein.** Cells were mock-transfected (Mock) transfected with pCAGGS/S-FL (S-FL) and stained using (i) PD4, (ii) PD5, (iii) sc23, (iv) sc29 and(iv) PolyS as indicated, and imaged using immunofluorescence (IF) microscopy (objective ×20 magnification).
